# Plant-associate interactions and diversification across trophic levels

**DOI:** 10.1101/2021.07.29.454357

**Authors:** Jeremy B. Yoder, Albert Dang, Caitlin MacGregor, Mikhail Plaza

## Abstract

Interactions between species are widely understood to have promoted the diversification of life on Earth, but how interactions spur the formation of new species remains unclear. Interacting species often become locally adapted to each other, but they may also be subject to shared dispersal limitations and environmental conditions. Moreover, theory predicts that different kinds of interactions have different effects on diversification. To better understand how species interactions promote diversification, we compiled population genetic studies of host plants and intimately associated herbivores, parasites, and mutualists. We used Bayesian multiple regressions and the BEDASSLE modeling framework to test whether host and associate population structures were correlated over and above the potentially confounding effects of geography and shared environmental variation. We found that associates’ population structure often paralleled their hosts’ population structure, and that this effect is robust to accounting for geographic distance and climate. Associate genetic structure was significantly explained by plant genetic structure somewhat more often in antagonistic interactions than in mutualistic ones. This aligns with a key prediction of coevolutionary theory, that antagonistic interactions promote diversity through local adaptation of antagonists to hosts, while mutualistic interactions more often promote diversity via the effect of hosts’ geographic distribution on mutualists’ dispersal.

---

**I**nteractions between species have long been recognized as key drivers in the diversification of life on Earth (Agrawal and Zhang, 2021; Darwin, 1859; Ehrlich and Raven, 1964; Farrell et al.,1992; Grant, 1949; Hembry et al., 2014; Thompson, 2005). Intimate interactions in particular — those in which a parasite or mutualist spends much of its life in association with a single host individual — are implicated in elevated rates of diversification (Cruaud et al., 2012; Farrell, 1998; Futuyma and Agrawal, 2009; McKenna et al., 2019; Mitter et al., 1988), patterns of phylogenetic congruence between host and associate lineages (Althoff et al., 2012; Cruaud et al., 2012; Escudero, 2015; Liu et al., 2013), and host-associated differentiation within species (Althoff, 2008; Drès and Mallet, 2002; Peterson and Denno, 1998; Schneider et al., 2016; Stireman et al., 2006). However, it remains unclear how often selection created by interacting species directly contributes to the formation of reproductive isolation, and there is a building consensus that different forms of interaction have different effects on diversification.

There are multiple processes by which species interactions may promote diversification, operating at time scales ranging from a few growing seasons to millions of years (Agrawal and Zhang, 2021; De Vienne et al., 2013; Janz, 2011; Thompson, 2005). The classic escape-and-radiate model predicts cycles of alternating diversification — first in the hosts or victims, then in the associates or enemies — driven by the evolution of defenses and counter-defenses (Ehrlich and Raven, 1964; Futuyma and Agrawal, 2009; Janz, 2011). Escape-and-radiate processes should result in associated clades within larger interacting lineages, such as Ehrlich and Raven’s butterflies and their larval host plants 1964, but not necessarily congruence at lower levels of biological organization. This is because the diversification occurs asynchronously — a victim clade diversifies after “escaping” the association with the help of a new defense, then the antagonist clade diversifies after overcoming that defense (Thompson, 2005).

At a smaller scale, intimately interacting species have also been predicted to show patterns of contemporaneous speciation (Forbes et al., 2009). Adaptation to an interacting species has been widely shown to create local adaptation and the beginnings of ecological speciation (Alstad 1998; Capelle and Neema 2005; Hanks and Denno 1994; Laine 2005; reviewed by Hargreaves et al. 2020; Hoeksema and Forde 2008; Runquist et al. 2020). However, evidence for speciation directly attributable to particular interactions has been surprisingly sparse (Althoff et al., 2014; Hembry et al., 2014). Within species, local adaptation driven by a species interaction may often be detectable as *ecological isolation*, in which the population structure of one interacting species parallels that of the other (Nosil et al. 2003; Peterson and Denno 1998; Wang and Bradburd 2014; Fig. 1). Such correlated population structure is not proof positive of coevolution with an interacting species, though — it may also evolve if both species have similar dispersal limitations (Wright, 1943) or both become locally adapted to variation in their shared environments (Futuyma and Peterson, 1985; Nuismer et al., 2010). Conversely, if alleles that determine hosts’ matching to or resistance against the associate species are distributed among populations in a manner that deviates from overall host population structure — a potential result of local adaptation, especially for a strongly selected trait determined by one or a few loci — coevolution between the two species could conceivably make their population structures *less* similar. Therefore, failure to find correlated population structures in a host plant and an associate species does not eliminate the possibility that one exerts selection on the other.

**Fig. 1.**
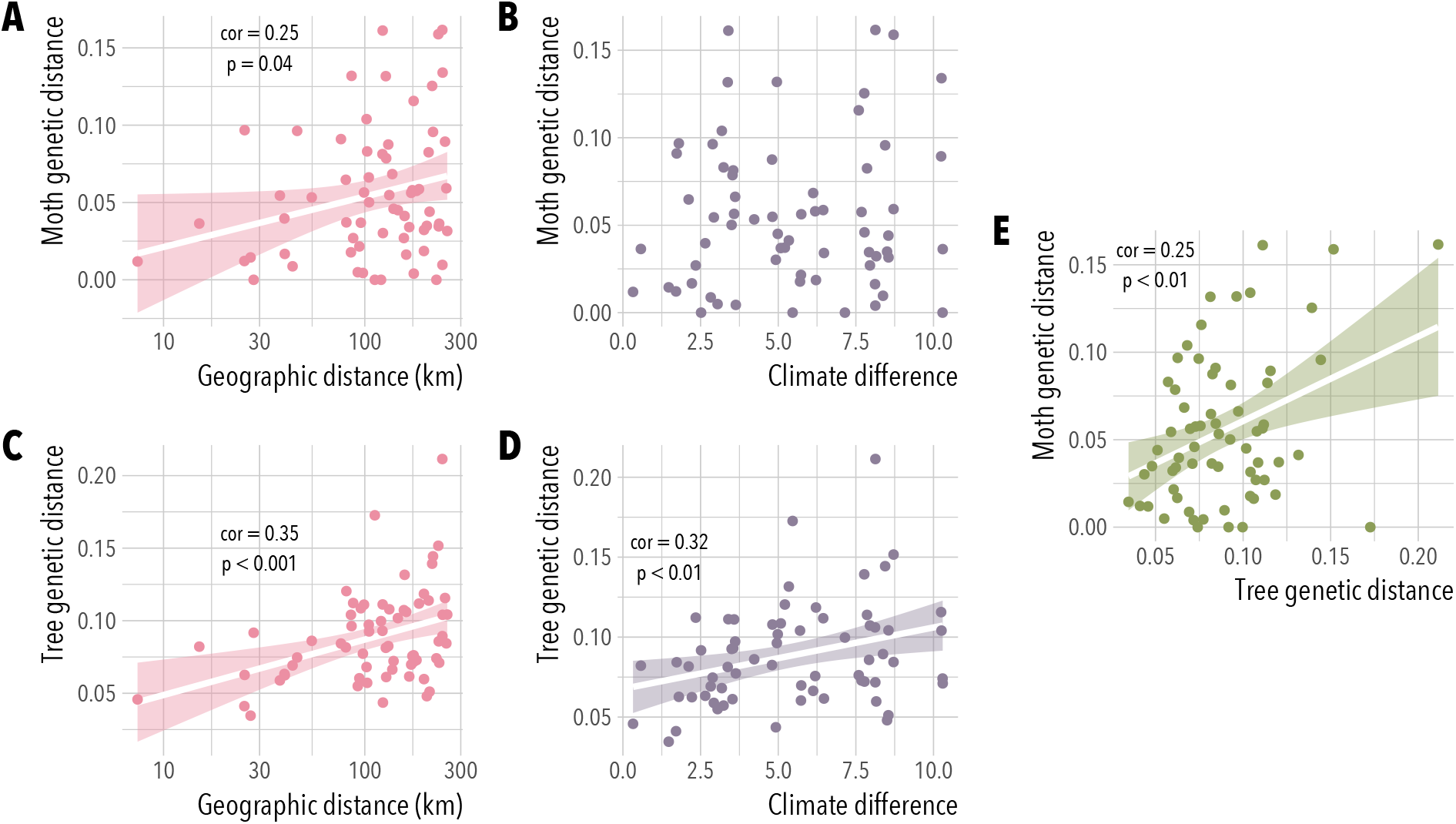
An example with data from western Joshua tree (*Yucca brevifolia*) and the pollinating, seed-feeding yucca moth *Tegeticula synthetica* (Yoder et al., 2013). Moth genetic distances (*F_ST_* /(1 − *F_ST_*)) are significantly correlated with geographic distance (A; Spearman’s *ρ* = 0.25, *p* = 0.04) but not with climate differences between sites (B); Joshua tree genetic distances show significant correlations with distance (C; *ρ* = 0.35, *p* < 0.001) and with climate (D; *ρ* = 0.32, *p* < 0.01). A significant positive correlation between Joshua tree genetic distance and moth genetic distance (E; *ρ* = 0.35, *p* < 0.05) may be consistent with local adaptation to the trees creating ecological isolation in the moths.

Furthermore, the nature of a species interaction probably determines whether or not it creates ecological isolation (Yoder, 2016). Antagonistic interactions such as predation or parasitism can create local arms races and cycles of allele frequencies or interacting phenotypes (Gomulkiewicz et al., 2000; Nuismer, 2006; Nuismer et al., 2010; Ridenhour and Nuismer, 2007; Yoder and Nuismer, 2010). These processes are expected to lead to local adaptation between populations of interacting species, and potentially co-diversification. In contrast, mutualistic interactions may often generate species-wide stabilizing selection, if the mutualists benefit from matching whatever traits mediate the interaction (Kiester et al., 1984; Yoder and Nuismer, 2010). In this case, correlated population structures would be more likely to arise via isolation by distance or local adaptation to shared environments (Nuismer et al., 2010), and theory in multiple frameworks has found that mutually beneficial interactions tend to promote less divergence or diversification than antagonistic interactions, under otherwise comparable circumstances (Kopp and Gavrilets, 2006; Maliet et al., 2020; Yoder and Nuismer, 2010).

Interactions between plants and the parasites, symbionts, and herbivores they host provide a potentially useful venue to examine congruence in the population structure of intimately interacting species, and to test contrasting predictions about the outcomes of different interaction types. Because they are a fundamental resource, plants are a substrate to which associated species must adapt, and as a limiting factor that may shape associates’ biogeography (Futuyma and Peterson, 1985). Plants’ impact on their associates’ local adaptation and population genetic structure has been a subject of study for decades (Blakley, 1982; Drès and Mallet, 2002; Futuyma and Peterson, 1985; Matsubayashi et al., 2010; Peterson and Denno, 1998). However, most population genetic studies in this literature examine only the plant, or the associate — or focus on adaptation of associates to contrasting host species rather than adaptation of associate populations to host populations (or vice versa) (Drès and Mallet, 2002; Futuyma and Peterson, 1985; Matsubayashi et al., 2010; Rausher, 1983). These trends reflect the need to simplify study design and interpretation, but they mean that we may miss intraspecific selection dynamics that are the ultimate cause of larger-scale evolutionary patterns (Thompson, 2005, 2013). Fortunately, studies of within-species population genetic data for plants and their associates are becoming both more practicable and more common. Compilation and synthesis of such studies can reveal patterns about the evolutionary importance of associates’ local adaptation to host plants, and may also guide future work in this line of inquiry by revealing biases and gaps in the range of study systems examined so far.

Here, we examine the congruence of population genetic structure in plants and associated species across a data set compiled from published studies. We test the hypothesis that host plants’ population structure predicts the population structure of associate species using meta-analysis of correlations between genetic, geographic and environmental distances, then use Bayesian multiple linear regression and a modeling framework specifically designed to study ecological isolation to examine the effect of host plant population structure on their associates’ genetics, over and above confounding effects of geography and climate. We find evidence that associates’ population structure often parallels their host plants’ population structure, in both mutualistic and antagonistic interactions, though this effect is somewhat stronger in antagonistic interactions than in mutualistic ones. We discuss these results in light of coevolutionary theory, and suggest future directions for population genetic studies of plant-associate interactions as robust genetic data becomes more accessible for non-model organisms.

## Methods

### Literature search

We compiled papers reporting population genetic data for plants and associated species from our personal collections, and using the Google Scholar search engine (scholar.google.com). We searched varying combinations of keywords referring to population structure (*structure*, *Fst*, or *"pairwise Fst"*), keywords referring to species interactions (*mutualism*, *pathogen*, *parasite*, *symbiont*, *pollinator*, *herbivore*, or *coevolution*), and always included the keyword *plant*. All keyword sets returned at least 1,500 results, and many had more than 20,000; we reviewed titles and abstracts for the first 200 papers returned in each search, and generally found that relevant results were not found after the first 100. We found, in total, 24 papers discussing or reporting population genetic data for both a plant species and at least one associated mutualist, herbivore, or pathogen. Of these, we retained papers that provided either pairwise genetic differentiation estimates, or genetic data that would support such estimates, for both a host plant and an associate species at four or more geographically distinct sites. As a check on the thoroughness of our literature search, we used Google Scholar to examine all papers citing the final set of studies. The final compiled data set included results from 15 published papers (Table 1).

**Table 1.** Plant-associate interactions in our compiled data set, the broad type of each associate, the number of sampling sites with data for both species, and the type of genetic markers used for each

| Plant/ Associate | Associate type | Sites | Markers for |  | Source |
| --- | --- | --- | --- | --- | --- |
|  |  |  | Plant | Associate |  |
| *Camellia japonica*/ *Curculio camelliae* | antagonist | 6 | SSR | sequence | Toju et al. (2011)* |
| *Datura stramonium*/ *Lema trilineata* | antagonist | 4 | SSR | allozyme | Garrido et al. (2012)* |
| *Ficus hirta*/ *Valisia* sp. | mutualist | 30 | SSR | SSR | Yu et al. (2019) |
| *Ficus pumila*/ *Wiebesia* sp. | mutualist | 24 | SSR | SSR | Liu et al. (2013) |
| *Hakea recurva*/ *Ameyma gibberula* | antagonist | 8 | SNP | SNP | Walters et al. (2021) |
| *Hirtella phyophora*/ *Allmoerus decemarticulatus* | mutualist | 14 | SSR | SSR | Malé et al. (2016) |
| *Lysimachia vulgaris*/ *Macropis europaea* | mutualist | 45 | AFLP | AFLP | Triponez et al. (2015) |
| *L. vulgaris*/ *M. fulvipes* | mutualist | 17 | AFLP | AFLP | Triponez et al. (2015) |
| *Medicago lupulina*/ *Ensifer* sp. | mutualist | 28 | SNP | sequence | Harrison et al. (2017) |
| *Pinus banksiana*/ *Arceuthobium americanum* | antagonist | 11 | AFLP | AFLP | Jerome and Ford (2002)*† |
| *P. contorta*/ *A. americanum* | antagonist | 10 | AFLP | AFLP | Jerome and Ford (2002)*† |
| *Populus angustifolia*/ *Aceria parapopuli* | antagonist | 10 | SSR | sequence | Evans et al. (2013)* |
| *Rhus chinensis*/ *Schlectendalia chinensis* | antagonist | 8 | SSR | SSR | Ren et al. (2007)*† |
| *Roridula gorgonius*/ *Pameridea roridulae* | mutualist | 14 | allozyme | allozyme | Anderson et al. (2004)*† |
| *Silene latifolia*/ *Hadena bicruris* | mutualist | 9 | SSR | SSR | Magalhaes et al. (2011) |
| *Vachellia drepanolobium*/ *Crematogaster mimosae* | mutualist | 5 | SNP | SNP | Boyle et al. (2019) |
| *V. drepanolobium*/ *Crematogaster nigriceps* | mutualist | 7 | SNP | SNP | Boyle et al. (2019) |
| *V. drepanolobium*/ *Tetraponera penzigi* | mutualist | 9 | SNP | SNP | Boyle et al. (2019) |
| *Yucca brevifolia*/ *Tegeticula synthetica* | mutualist | 13 | SSR | SSR | Yoder et al. (2013) |
| *Y. jaegeriana*/ *T. antithetica* | mutualist | 10 | SSR | SSR | Yoder et al. (2013) |

\*Provided genetic distances but not original genotype data
†Insufficient information to derive climate data for sampling sites

### Data compilation

We assembled our working data set and conducted all analyses in R (version 4.0.0, R Core Team 2020). The final working data set included data from 20 plant-associate pairs, reported in 16 different published studies (Table 1). The type of genetic marker used varied by study and even between plants and associates in the same study (Table 1), reflecting the challenge, until recently, of obtaining similar-quality genetic data for two interacting non-model species. Simple sequence repeat markers (SSRs, or microsatellites) were the most common marker type, with amplified fragment length polymorphisms (AFLPs) and allozyme polymorphisms represented in multiple cases as well. For analysis we required pairwise genetic, climatic, and geographic distances for both the host plant and one or more associated species, across the same sampling sites. Most of the papers in our final set reported these values directly, either in the main text or as supplementary information. However, to minimize variation in the methods deriving genetic, geographic, and climatic distances, we re-derived genetic distances for 13 plant-associate pairs for which we could obtain original genotype data, and we re-estimated climatic distances in all cases where we could determine the locations of sampling sites with sufficient precision from information provided (Table 1).

#### Genetic distances

Because we used the population genetic modeling framework BEDASSLE (Bradburd et al. 2013) for downstream analysis (see below), we calculated *F_ST_* between pairs of sampling sites using the `calculate.all.pairwise.Fst()` function provided in the BEDASSLE package, which follows Weir and Hill’s (2002) method. We reformatted genotype data to conform to meet BEDASSLE’s requirement for biallelic loci; in the case of SNP loci with more than two alleles, and all SSR and fragment polymorphism data, we followed a recommended recoding scheme to treat alternate alleles as occurring at separately coded (pseudo)loci (G. Bradburd, *pers. comm*.). For instance, a locus with alleles A, B, and C, would be recoded as three loci, the first with alleles A/not A, the second with alleles B/not B, and the third with alleles C/not C. Six studies representing seven plant-associate pairs (Anderson et al. 2004; Garrido et al. 2012; Jerome and Ford 2002; Magalhaes et al. 2011; Ren et al. 2007; Toju et al. 2011; Table 1) provided genetic distances between sites but not genotype data we could use to estimate *F_ST_*; in these cases we used reported genetic distances.

#### Geographic distances

Where papers or supporting information provided sampling site locations, either as latitude and longitude or as maps with sufficient detail to determine latitude and longitude, we used the site locations to calculate great-circle distances between sampling sites, in kilometers, with the `rdist.earth()` function provided in the `fields` package (Nychka et al. 2017). Four studies representing five plant-associate pairs (Anderson et al. 2004; Jerome and Ford 2002; Magalhaes et al. 2011; Ren et al. 2007; Table 1) provided geographic distances between sampling sites but insufficient information to infer sampling site locations; in these cases we used reported geographic distances.

#### Climatic distances

We extracted Bioclim climate data (Woldclim version 2.1, averages from 1970-2000; Fick and Hijmans 2017; Hijmans 2020) for sampling site locations. We then calculated climate differences between sampling sites by performing principal components analysis on site climate values and calculating between-site Euclidean distances in PCA space, using the base R functions `prcomp()` and `dist()`. The PCA transformation should account for the fact that the 19 Bioclim variables are strongly intercorrelated, by reorienting the data along major axes of variation, while at the same time allowing us to avoid the difficulty of selecting specific climate variables most relevant to each (or all) species pair(s). In the case of plant-associate pairs from studies that provided insufficient information to infer sampling site locations (Table 1), we performed analyses without climate distance as a variable.

### Analyses

To describe relationships among genetic, environmental (climatic), and geographic distances, we first calculated simple correlations between all possible pairwise combinations of distance metrics for each plant-associate pair, as Spearman rank correlations. We used the base-10 logarithm of geographic distances, and the standard transformation of genetic distances as *D_genetic_*/(1 − *D_genetic_*) (Rousset, 1997). To estimate confidence intervals around pairwise correlations, we bootstrapped the distance data, using the `bootstraps()` function of the `rsample` package (Kuhn et al., 2020), with 1000 bootstrap permutations. We also used correlation coefficient estimates to check for publication bias, by testing for a correlation between individual studies’ sample sizes (as total number of pairwise comparisons) with the estimated correlation coefficients for isolation by geography and isolation by climate in the host and associate, as well as the host-associate correlation. A significant negative correlation between sample size and effect is consistent with the “file drawer” bias, in which studies are less likely to be published if they fail to find a hypothesized effect (Nakagawa et al., 2017).

To test whether host plant population structure creates ecological isolation for associates while accounting for shared geography and environmental factors, we fitted multiple linear regressions predicting associate genetic distances with additive linear effects of geographic distance, climate distance (if available), and host plant genetic distance. To cope with the highly non-normal distribution of the data, we fitted models using the Bayesian framework implemented in the brms package (Bürkner, 2018, 2017), with the response modeled as a zero-inflated beta distribution. To account for the spatial non-independence of between-site distance metrics, we assessed whether regression coefficients were significantly different from zero following the approach of multiple matrix regression with randomization (Wang 2013), generating a null distribution by permuting the response matrix (i.e., associate genetic distances) and refitting the Bayesian multiple linear regression. We identified a regression coefficient as significantly different from zero if the 95% density interval of the corresponding coefficient in 1000 permuted regressions did not contain zero.

For the 13 plant-associate pairs with genotype data available (Table 1), we further explored isolation by distance, climate, and host plant using the modeling framework BEDASSLE, which is explicitly designed to estimate and compare the ecological isolation attributable to different environmental factors (Bradburd et al. 2013). For each plant-associate pair, we re-coded genotype data to conform to BEDASSLE’s requirements (see data compilation, above) and estimated, for the associate species, the isolating effects (*alpha* terms, in the BEDASSLE model) of geographic distance, climate distances between sampling sites, and host genetic distances between sampling sites. Following recommended practice, we performed pilot runs of the BEDASSLE Markov chain Monte Carlo model-fitting procedure and inspected parameter estimate and acceptance rate traces, adjusting the size of parameter changes allowed between MCMC steps until the acceptance rates showed the Markov chain achieving stationarity. When we had identified appropriate parameter change values, we ran a final model-fitting procedure for 5 × 10^6^ iterations. We used the last 1 × 10^6^ generations for posterior parameter estimation and discarded the rest as burnin.

Finally, we used formal meta-analysis to examine the overall strength of the pairwise distance correlations, predictor effects in the multiple linear regressions, and ecological isolation estimates from BEDASSLE. We used the random-effects methods for correlations (the `metacor()` function) and mean effect estimates (the `metamean()` function) provided in the `meta` package (Balduzzi et al., 2019). To test for differences between antagonistic and mutualistic interactions, we included interaction types as a grouping variable in each random-effects meta-analysis. Associate taxa proved to be widely and unevenly phylogenetically dispersed, including a bacterium, two parasitic plants, and multiple insects, but also with three cases in which congeners or the same species were represented in multiple pairs (Table 1: *Arceuthobium*, *Crematogaster*, *Macropis*, and *Tegeticula*). Because of the highly skewed range of phylogenetic distances among associates, we did not employ a formal control for phylogenetic non-independence in our meta-analyses, but we tested the robustness of our conclusions by systematically re-running each meta-analysis with each possible permutation of the data set created by excluding one member from each of these three pairs of close relatives.

## Results

Our literature search obtained 15 papers reporting population genetic data from 20 plant-associate pairs (Table 1). Seven of the 20 pairs had antagonistic interactions, with the associates being herbivores or parasites, and the others were mutualistic interactions such as pollination, protection, or nutrient exchange. In 16 pairs, the associate was an insect; three pairs consisted of parasitic plants, *Arceuthobium americanum* on two different conifer hosts and *Amyema gibberula* on its host *Hakea recurva*; and the final pair consisted of nitrogen-fixing rhizobial bacteria, genus *Ensifer*, and their legume host *Medicago lupulina*. In addition to the two data sets with *A. americanum* as the associate, cuckoo bees (genus *Macropis*), plant-ants (genus *Crematogaster*), and yucca moths (genus *Tegeticula*) were each represented in two host-associate pairs. Of the 15 original papers providing data that we could use for synthesis, eight included explicit comparisons of host plant and associate population structure, generally a regression of associate genetic distances on host plant genetic distances (Anderson et al. 2004; Evans et al. 2013; Harrison et al. 2017; Jerome and Ford 2002; Liu et al. 2013; Magalhaes et al. 2011; Ren et al. 2007; Triponez et al. 2015).

The spatial scale of the compiled data sets varied over orders of magnitude (Table 2), with median geographic distance between sampled sites in each study ranging from less than 16km (in the *Camellia japonica*/*Curculio camelliae* pair; Toju et al. 2011) to more than 1,000 km (*Lysimachia vulgaris*/*Macropis fulvipes*; Triponez et al. 2015). Genetic distances were also quite variable, with median *F_ST_* for host plants as low as 0.04 (*Pinus banksiana*; Jerome and Ford 2002) and as high as 0.9 (*Roridula goronius*; Anderson et al. 2004, and median *F_ST_* for associates ranging from 0.03 (*Tegeticula antithetica*; Yoder et al. 2013) to 0.6 (*Tetraponera penzigi*; Boyle et al. 2019). (Genetic distances for one species pair, *Camellia japonica* and *Curculio camel-liae*, were provided as Nei’s *D*, which scales differently than *F_ST_*; Toju et al. 2011.) This variation is perhaps not surprising given the range of taxonomy represented in the compiled dataset, and we generally assume that the authors of the original studies designed their sampling schemes to best capture population structure of their study organisms based on those species’ dispersal capabilities and mating systems.

**Table 2.** Geographic and genetic distances covered by the data compiled for each plant-associate pair

| Plant/ Associate | Geographic dist (km)* | Genetic dist, plant*† | Genetic dist, associate*† |
| --- | --- | --- | --- |
| *Camellia japonica*/*Curculio camelliae* | 15.9 (2.6, 19.5) | 0.20 (0.15, 0.53) | 0.50 (0.00, 2.10) |
| *Datura stramonium*/*Lema trilineata* | 132.4 (54.6, 241.0) | 0.34 (0.25, 0.55) | 0.10 (0.03, 0.16) |
| *Ficus hirta*/*Valisia* spp | 694.6 (100.0, 1998.4) | 0.11 (0.03, 0.27) | 0.41 (0.05, 0.55) |
| *F. pumila*/*Wiebesia* sp. 1 | 27.2 (4.7, 172.0) | 0.09 (0.02, 0.23) | 0.10 (0.02, 0.21) |
| *Hakea recurva*/*Amyema gibberula* | 151.9 (62.2, 328.9) | 0.08 (0.04, 0.22) | 0.44 (0.11, 0.67) |
| *Hirtella physophora*/*Allmoerus decemarticulatus* | 54.8 (14.6, 75.0) | 0.05 (0.02, 0.11) | 0.19 (0.08, 0.26) |
| *Lysimachia vulgaris*/*Macropis europaea* | 610.9 (121.6, 1358.1) | 0.16 (0.01, 0.48) | 0.06 (0.00, 0.42) |
| *L. vulgaris*/*M. fulvipes* | 1034.3 (114.1, 2349.1) | 0.28 (0.05, 0.57) | 0.12 (0.00, 0.53) |
| *Medicago lupulina*/*Ensifer medicae*, *E. meliloti* | 131.6 (18.5, 459.0) | 0.36 (0.12, 0.79) | 0.11 (0.00, 0.71) |
| *Pinus banksiana*/*Arceuthobium americanum* | 14.7 (7.3, 22.7) | 0.04 (0.00, 0.10) | 0.21 (0.11, 0.33) |
| *P. contorta*/*A. americanum* | 18.2 (12.1, 23.9) | 0.19 (0.07, 0.28) | 0.08 (0.02, 0.20) |
| *Populus angustifolia*/*Aceria parapopuli* | 454.2 (135.9, 901.7) | 0.10 (0.01, 0.40) | 0.13 (0.00, 0.85) |
| *Rhus chinensis*/*Schlectendalia chinensis* | 123.0 (36.6, 241.8) | 0.19 (0.08, 0.26) | 0.23 (0.18, 0.29) |
| *Roridula gorgonius*/*Pameridea roridulae* | 23.5 (0.5, 119.6) | 0.90 (0.00, 0.97) | 0.05 (0.00, 0.68) |
| *Silene latifolia*/*Hadena bicruris* | 312.3 (14.6, 592.5) | 0.06 (0.02, 0.12) | 0.03 (0.01, 0.06) |
| *Vachellia drepanolobium*/*Crematogaster mimosae* | 60.8 (11.8, 146.0) | 0.16 (0.07, 0.24) | 0.20 (0.05, 0.29) |
| *V. drepanolobium*/*C. nigriceps* | 93.1 (12.5, 152.3) | 0.22 (0.07, 0.41) | 0.50 (0.01, 0.73) |
| *V. drepanolobium*/*Tetraponera penzigi* | 71.4 (10.6, 149.0) | 0.22 (0.06, 0.41) | 0.60 (0.16, 0.82) |
| *Yucca brevifolia*/*Tegeticula synthetica* | 127.4 (21.4, 251.8) | 0.08 (0.04, 0.14) | 0.04 (0.00, 0.14) |
| *Y. jaegeriana*/*T. antithetica* | 106.7 (22.3, 245.6) | 0.08 (0.03, 0.14) | 0.03 (0.00, 0.08) |

\*Median (95% density interval)
†Genetic distances are $F_{ST}$ except for *Camellia japonica*/*Curculio camelliae*, which use Nei's D (Toju et al., 2011).

### Correlations with genetic distances

Pairwise correlations between genetic and geographic distance (i.e., isolation by distance), between genetic and climate distance (isolation by climate), and between genetic distances for plants and associate species (isolation by host) varied considerably across plant-associate pairs (Fig. 2). Host plants showed significant isolation by distance (Spearman rank correlation, *ρ* > 0 and bootstrapped *p* < 0.05) in 11 of 20 pairs, and significant isolation by climate differences in 7 of 16 pairs for which climate data were available. Associates showed correlations consistent with isolation by geographic distance in 14 of 20 pairs, and isolation by climate in 10 of 16 pairs.

**Fig. 2.**
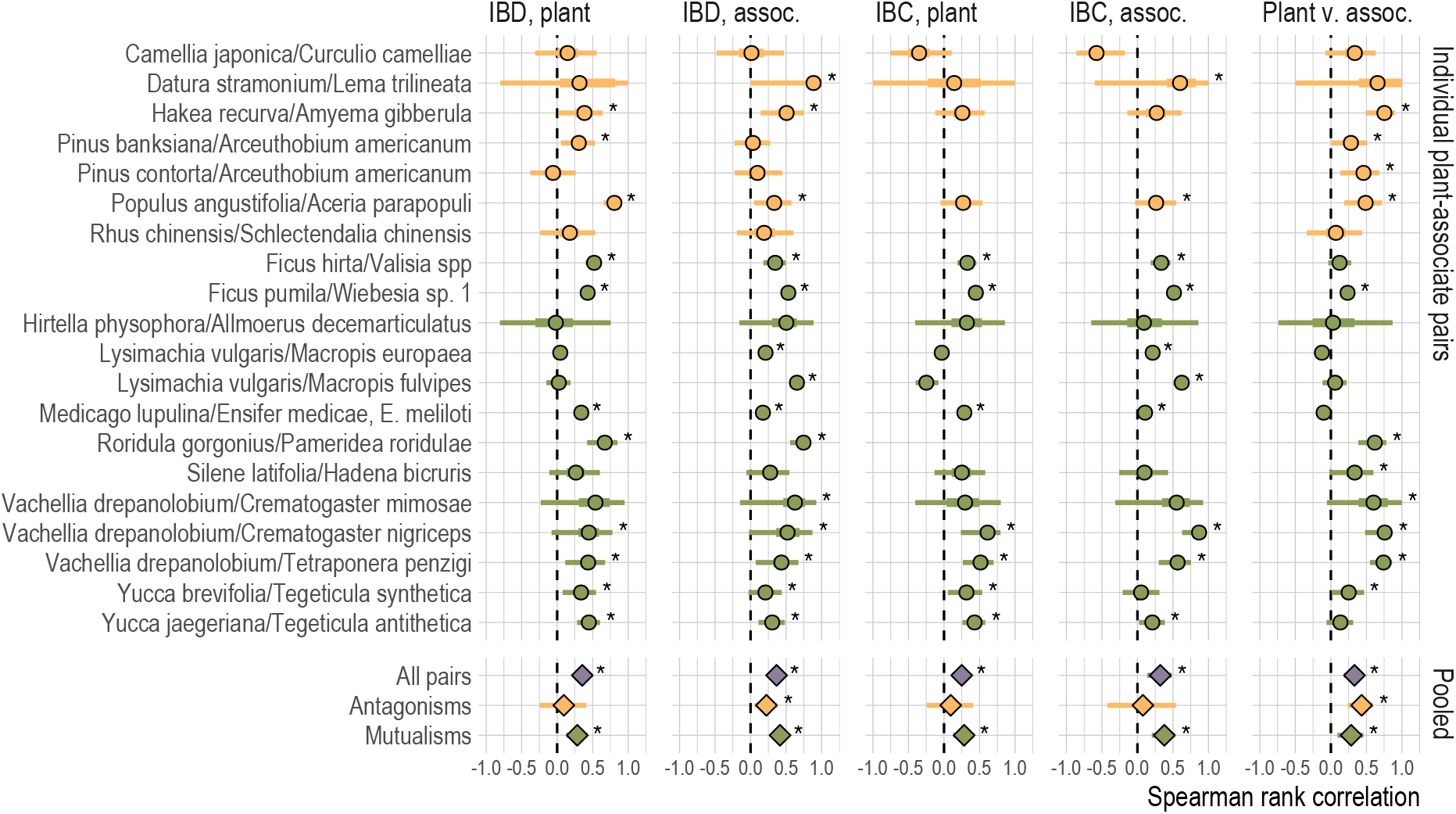
Correlations between genetic, geographic, and climate distances in the plant-associate data sets we compiled, for each of 20 individual plant-associate pairs and for meta-analytic summaries across mutualisms, antagonisms, and all pairs. For individual pairs, points indicate Spearman rank-sum correlation *ρ*, bars give 95% (thin) and 50% confidence (thick) intervals based on 1000 bootstrap replicates. For meta-analytic pooled estimates, points give estimated mean and bars give 95% confidence intervals. Asterisks indicate correlations greater than zero with bootstrap *p* < 0.05. Antagonistic interactions are colored yellow, mutualistic ones green.

Plant and associate genetic distance were significantly correlated, potentially consistent with associates experiencing isolation by host plant, in 11 of 20 plant-associate pairs — in 4 of the 7 antagonistic interactions, and 7 of the 13 mutualistic interactions. For two pairs we found a significantly nonzero *negative* correlation between host and associate genetic distances; this pattern, which indicates that associates interacting with more closely related host populations are themselves more genetically differentiated, is also reported in the original study of one of the pairs, the loosestrife *Lysimachia vulgaris* and the pollinating bee *Macropis europaea*, by Triponez et al. (2015). Across the 20 plant-associate pairs, significant correlations between plant and associate genetic structure were not consistently seen in conjunction with isolation by distance or isolation by climate in either plants or associates.

The general signal revealed by meta-analysis was also that plant and associate genetic distances are significantly and positively correlated (Fig. 2; pooled estimate of Spearman’s *ρ* = 0.33, 95% confidence interval from 0.19 to 0.46). The meta-analytic pooled correlation between plant and associate genetic distances was also significantly greater than zero within antagonistic and mutualistic interactions (for antagonistic interactions, pooled *ρ* = 0.43, 95% CI 0.24 to 0.60; for mutualistic, *ρ* = 0.29, 95% CI 0.09 to 0.46). Although the mean correlation was somewhat greater for antagonistic interactions than for mutualistic interactions, this difference was not greater than expected by chance (*p* = 0.27 for group differences in the random-effects meta-analysis). Excluding closely related associates from the data set did not qualitatively alter any of these results. Overall, this is consistent with associates experiencing widespread isolation by host plant, across different types of interactions.

However, meta-analysis also found correlations with distance and climate that may be confounded with the correlations in genetic distances see above. Meta-analysis across all plant-associate pairs found significant isolation by distance for plants and associates (pooled *ρ* = 0.35, 95% CI 0.23 to 0.46 for plants; *ρ* = 0.37, 95% CI 0.26 to 0.46 for associates) and significant isolation by climate for plants and associates (*ρ* = 0.25, 95% CI 0.11 to 0.38 for plants; *ρ* = 0.32, 95% CI 0.14 to 0.48 for associates). These results were generally robust to pruning closely related associates.

Meta-analysis of only antagonistic interactions or only mutualistic interactions found similar results to the overall pattern, with the exception that pooled correlations between genetic distance and climate distance were not greater than zero in antagonistic interactions, potentially consistent with a smaller or less consistent effect of isolation by climate in these systems (*ρ* = 0.09, 95% CI −0.24 to 0.41 for plants; *ρ* = 0.08, 95% CI −0.42 to 0.54 for associates). These differences between interaction types were not greater than expected by chance in the random effects model, however (*p* > 0.05 for both isolation by distance and isolation by climate). These results were also robust to pruning closely related associates, with the exception that pruning the *Pinus banksiana*/*Arceuthobium americanum* species pair resulted in non-significant pooled isolation by distance correlations for host plants in antagonistic interactions.

Comparing the pairwise correlation coefficients to the sample sizes for individual studies revealed no pattern consistent with publication bias in isolation by distance for host or associate taxa, or in isolation by climate (Spearman rank tests for correlation between study sample size and the respective correlation coefficients; *p* > 0.2 in all cases). However, we did find a significant negative correlation between sample size and the correlation coefficients for plant and associate population structure (*ρ* = −0.60, *p* = 0.005). This pattern was also significant when assessed only for mutualistic species pairs (*ρ* = −0.63, *p* = 0.02), though not for antagonistic pairs (*ρ* = −0.29, *p* = 0.53). This is consistent with the prospect that studies failing to find correlated population structure in host plants and associated taxa are more likely to go unpublished.

### Multiple linear regressions

Bayesian multiple linear regressions explaining associate population structure by the joint effects of geographic distance, climate (where available), and host plant genetic distance returned results that generally paralleled patterns in the simple pairwise correlations (Fig. 3). Correlations between plant and associate genetic distances strongly predicted multiple regression estimates of the effect of plant genetic distance on associate genetic distance (Spearman’s *ρ* = 0.60, p = 0.005; Figs 2,3). However, multiple regression models included a significantly nonzero effect of isolation by host plant in only eight plant-associate pairs (40% of the 20; Fig. 3). In general, significant effects of isolation by climate and geography were less common than significant effect estimates for isolation by host plant; none of the antagonistic associates showed significant isolation by distance, and only one, the weevil *Curculio camelliae*, showed significant isolation by climate (Fig 3).

**Fig. 3.**
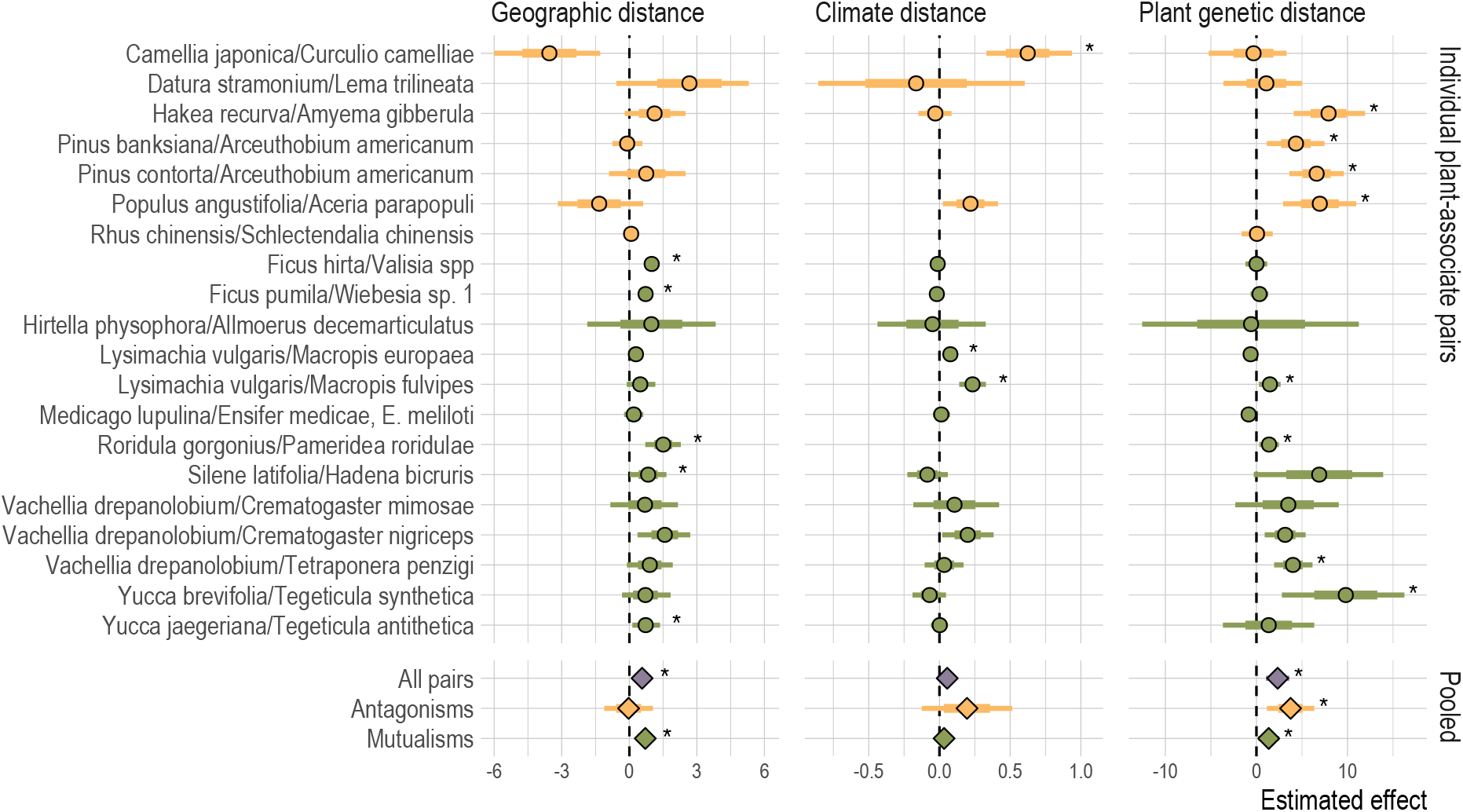
Estimated effects of geographic distance, climate distance, and host plant genetic distance on associate genetic distance in Bayesian multiple linear regressions for each of 20 individual plant-associate pairs; and meta-analytic summaries of the same effect estimates across mutualisms, antagonisms, and all pairs. Points indicate posterior mean effect, bars give 95% (thin) density intervals and standard error (thick). Asterisks indicate regression effects different from zero with permutational *p* < 0.05, and meta-analysis effect estimates for which the 95% confidence interval does not cross zero. Colors follow Fig. 2.

Meta-analysis across all plant-associate pairs found a significant effect of plant genetic distance (Fig. 3; estimate = 2.32, 95% CI 1.05 to 3.62) and a significant effect of geographic distance (estimate = 0.58, 95% CI 0.32 to 0.82); but the meta-analytic mean effect of climate was not significantly different from zero (estimate = 0.06, 95% CI −0.008 to 0.12). Pruning close relatives did not change these qualitative results. This is consistent with associates generally experiencing both isolation by distance and isolation by host plant, but not isolation by climate.

Meta-analysis of only antagonistic pairs found no significant isolation by distance (estimate = −0.03, 95% CI −1.12 to 0.51), but significant isolation by host plant (estimate = 3.75, 95% CI 1.15 to 6.35); whereas meta-analysis of the mutualist pairs found significant isolation by distance (estimate = 0.71, 95% CI 0.47 to 0.94) and significant isolation by host plant (estimate = 1.35, 95% CI 0.20 to 2.50). Isolation by climate was not significantly greater than zero in meta-analysis of either interaction type (estimate = 0.03, 95% CI −0.02 to 0.09 for mutualisms; estimate = 0.20, 95% CI −0.13 to 0.51 for antagonisms). The random-effects meta-analysis found that the difference in isolation by distance with associate type was not significantly greater than expected by chance (*p* = 0.19); and, although antagonisms showed greater mean isolation by host plant than mutualisms, this was also non-significant (*p* = 0.10). Pruning close relatives generally did not change the qualitative results — except that the pooled effect of isolation by host plant in mutualisms was not significantly different from zero when *Macropis fulvipes* was excluded, or when both *Macropis fulvipes* and *Crematogaster nigriceps* were excluded. Overall, this is consistent with antagonistic associates experiencing isolation by host plant but not by geographic distance, while mutualistic associates experience isolation by distance and, to a lesser extent, by host plant.

### Analysis with BEDASSLE

The 13 species pairs for which we could obtain genotype data produced widely varying estimates of isolating effects for geographic distance, climate distance, and host plant genetic differentiation in analysis with BEDASSLE (Fig. 4), and these isolating effects did not cleanly correlate with the equivalent pairwise correlation estimates or effects in multiple linear regression models (Spearman’s rank correlation of posterior median *α* terms versus correlation coefficients or effect estimates, *p* > 0.05 in all cases). All *α* terms for all species pairs had 95% posterior densities that did not cross zero, though this likely reflects a constraint of the BEDASSLE model, which enforces *α* terms greater than zero. Meta-analysis of *α* terms yielded more nuanced results: across all 13 species pairs, the pooled *α* terms for geographic distance and climate distance had 95% posterior density intervals that included zero (for geographic distance, median effect = 0.124, 95% density −0.025 to 0.274; for climate distance, median effect = 0.189, 95% density −0.002 to 0.379); while the pooled *α* term for host plant genetic distance was strongly significantly greater than zero (median effect = 1.041, 95% density 0.624 to 1.458), consistent with associate genetic differentiation explained predominately by host plant genetic differentiation.

**Fig. 4.**
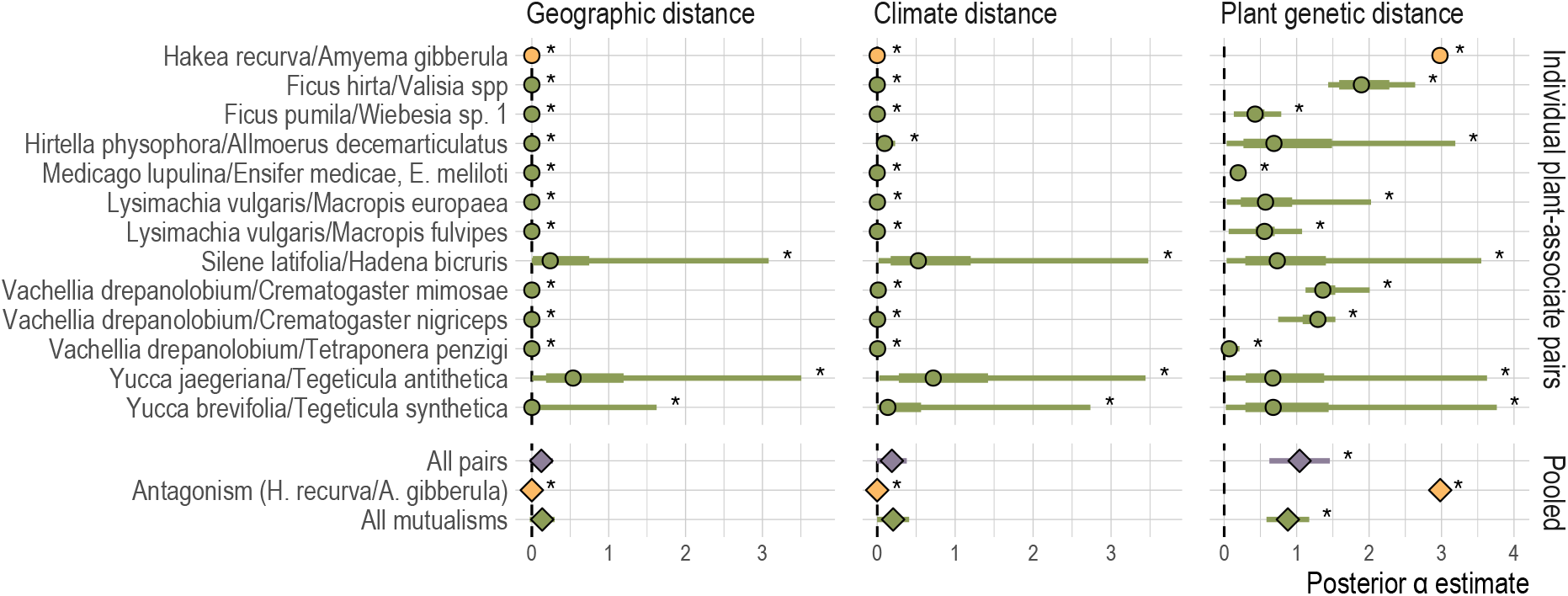
The isolating effects (*α*) of geographic distance, climate distance, and host plant genetic distance on associate species, as estimated in the BEDASSLE framework for each of 13 individual plant-associate pairs; and meta-analytic summaries of the same effect estimates across all pairs, across all mutualisms, and for the single antagonistic pair. For individual plant-associate pairs, points indicate posterior median effect, bars give 95% (thin) and 50% (thick) posterior density intervals; for meta-analytic pooled estimates, points indicate median pooled estimates, bars give 95% confidence intervals. Asterisks indicate posterior estimates and meta-analytic pooled estimates for which the 95% posterior density or 95% confidence interval does not cross zero. Colors follow Fig. 2

Only one of the 13 species pairs in the BEDASSLE analysis, the mistletoe *Amyema gibberula* and its host *Hakea recurva*, was antagonistic, sharply limiting our power to compare antagonistic and mutualistic interactions in meta-analysis. With this caveat, the random effects meta-analysis found that *Amyema gibberula* did not differ significantly from the mutualistic associates in posterior estimates of the *α* for geographic distance and climate distance; but antagonist had a posterior median *α* for host plant genetic distances substantially and significantly greater than the mutualists (Fig. 4; median pooled effect = 2.982, 95% posterior density from 2.981 to 2.983 interval for the antagonist; median = 0.879, 95% density 0.584 to 1.174 for the 12 mutualists; *p* < 0.0001 for the test of subgroup heterogeneity in the meta-analysis). Pruning close relatives from the meta-analysis did not change this qualitative result.

## Discussion

A wide range of experimental and observational evidence describes the ways in which plants may shape the evolutionary history of species that depend upon them. Population genetic data from plants and associates interacting across a shared landscape provides a look at local adaptation, the process by which the ecological dynamics of plant-associate interactions are translated into global patterns of biodiversity (Drès and Mallet, 2002; Futuyma and Peterson, 1985; Hembry et al., 2014; Peterson and Denno, 1998). Plants and their associates disperse and evolve across a shared landscape, and face the same geographically varying environments, even as associates adapt to their host plants (Futuyma and Peterson, 1985; Thompson, 2005); and different types of interactions may create differing plant-associate selective dynamics, and therefore different evolutionary outcomes (Maliet et al., 2020; Nuismer et al., 2010; Yoder, 2016).

Compiling published population genetic data for plant-associate pairs, we find evidence that host plants shape their intimate associates’ evolution through the effects of shared geographic distribution and through associates’ local adaptation to host plants — but generally not through adaptation to shared environmental conditions, specifically climate (Fig. 3). Pairwise correlations between genetic distances and geographic or climate distances are consistent with effects of isolation by distance and isolation by climate in both plants and associate species (Fig. 2). However, in multiple regressions accounting for the confounding among geography, climate, and host plant genetic distances, we see that climate differences have smaller impacts on associates’ genetic differentiation than either geographic distance or host plant genetic distances (Fig. 3). Accounting for geographic distance, climate, and plant genetic distance simultaneously in the multiple regression framework also reduces the number of cases in which we see significant correlations between plant population structure and the population structure of associate taxa. In the BEDASSLE framework, which is specifically designed to compare the population genetic effects of geographic and ecological isolation, we find that isolation attributable to host plants is stronger than that attributable to either geography or climate (Fig. 4).

The three analyses we employ here have different explanatory power. The pairwise correlations are purely descriptive, while the multiple regression and BEDASSLE analyses attempt to account for confounding among possible explanatory variables; and while BEDASSLE is designed specifically for studying ecological isolation with population genetic data, we are able to examine more cases using the multiple linear regression framework, given the data available. Overall, these results support a general pattern of associate species experiencing ecological isolation arising from the population genetic structure of their host plants — and this is consistent with associates locally adapting to their hosts.

It is possible that for some or all of the associate species we examine, ecological isolation may be driven by additional environmental factors that are not captured by the climate data we apply — anything from soil chemistry to other interacting species. The generally small effects of geographic distance estimated in the BEDASSLE analysis (Fig. 4) suggest that there is not another, unmeasured, spatially varying environmental factor that competes with host plant genetic structure. However, if the unmeasured factor is heavily co-linear with plant genetic structure (or, indeed, if plants experience ecological isolation due to that factor) we would expect to see essentially the same pattern that we find. Our results must therefore be understood as consistent with associates locally adapting to host plants, but not proof positive that they have done so.

The wide variation in spatial scales covered by the studies we have compiled and the range of genetic differentiation among the populations they sample (Table 2), raise the question of whether the same population genetic processes are captured in the 20 data sets we synthesize here. As noted above, we have assumed that the authors of the studies we have compiled planned their data collection with the natural history of their study species in mind, to best capture population structure and biologically relevant environmental variation. It is also true that a narrow majority of the original studies in our data set include an explicit comparison of host plant and associate population structure, as we set out to perform (eight of 15 studies: Anderson et al. 2004; Evans et al. 2013; Harrison et al. 2017; Jerome and Ford 2002; Liu et al. 2013; Magalhaes et al. 2011; Ren et al. 2007; Triponez et al. 2015) — these cases suggest that our use of the data is not inappropriate.

As with all synthesis of published research, our results may be shaped in part by biases in data collection or publication of studies fitting the criteria we require. All of the studies we compile here are cases in which the authors had some *a priori* expectation that the host plant and associate species influenced each other’s evolution at the population genetic level. It may seem somewhat less likely that studies conducted with this motivation would face the “file drawer” problem of being withheld from publication because of negative results, given the resources involved in collecting population genetic data for two species, and the possibility of finding meaningful patterns in the host if not the associate, or vice versa. However we do find a significant negative correlation between the strength of plant-associate population structure correlations and study sample size — so an important caveat to our analyses is that the data we have compiled may overestimate the general effect of associate species adapting to their host plants. Whether our comparison of antagonistic and mutualistic interactions is similarly biased is difficult to assess; on the one hand, we do not find a pattern consistent with publication bias in the data from antagonistic interactions, but on the other hand, the smaller number of studies representing antagonistic interactions makes the test for publication bias itself less powerful.

### Global patterns in plant-associate interactions

Overall, these results demonstrate that associates’ population structures often reflect those of their host plants, and that this pattern arises at least in part because of associates’ local adaptation to hosts (ecological isolation by host population) in addition to the effects of shared geography (isolation by distance). We find that the meta-analytic mean effect of climate differences on associate genetic distance is not significantly greater than zero, reflecting the fact that we found nonzero effects of climate in only three of 16 plant-associate pairs for which we could obtain climate data (Fig. 3). On the one hand, it may be surprising that adaptation to shared environments does not contribute to congruence in plant-associate population structure; but on the other it may be that climate differences are sufficiently conflated with geographic distance and host plant population structure that including this additional variable is not informative. Recent systematic reviews and meta-analysis have also found that local adaptation to abiotic factors is often weaker than local adaptation to biotic interactions (Hargreaves et al., 2020; Runquist et al., 2020), and the pattern we see here is consistent with these results.

Perhaps most interestingly, we find indications of different trends in different types of plant-associate interaction. In the multiple linear regression framework, we find a significantly nonzero effect of host plant genetic distances on the population structure of both mutualistic and antagonistic associates, but a somewhat stronger effect of host plant genetic distances for antagonistic associates (Fig. 3, meta-analysis within interaction types). This is recapitulated in our BEDASSLE analysis of a more limited set of plant-associate pairs, which finds a significantly stronger isolating effect of host plant genetic distances for the single antagonistic associate represented in the data set than for the 12 mutualists (Fig. 4). Finding stronger isolation by host plant for antagonistic associates aligns with the predictions of coevolutionary theory in multiple frameworks, which have found that antagonistic interactions mediated by phenotype differences or inverse matching can promote diversification of associates, whereas mutualistic interactions mediated by matching of host and associate traits tend to create stabilizing selection across populations (Kopp and Gavrilets, 2006; Maliet et al., 2020; Yoder and Nuismer, 2010). In the former case, the arms-race and inverse-frequency-dependent dynamics of antagonistic interactions should often mean that associates are adapted to their local host populations (Gomulkiewicz et al., 2000; Nuismer et al., 2007; Ridenhour and Nuismer, 2007), and therefore show isolation by host plant, as we find. In the latter case — mutualisms — we would expect that associates’ variation across populations would be driven more by drift and spatial isolation than by local adaptation to host plant populations, because hosts and associates would generally be selected to remain compatible across populations (Kiester et al., 1984; Yoder and Nuismer, 2010).

These expectations from theory are based on assumptions about the forms of selection operating in mutualistic versus antagonistic interactions, which have not been evaluated in most of the plant-associate pairs we examine here. However, educated inferences can be made many cases. For one antagonistic plant-associate pair in our data set *Camellia japonica*/*Curculio camelliae*, we have good documentation for arms-race dynamics mediated by physical defenses, the thickness of the host plant pericarp (Toju et al., 2011). In other plant-herbivore interactions such as *Datura stramonium*/*Lema trilineata* and *Populus angustifolia*/*Aceria parapopuli* we often see arms-race dynamics, with plants producing defensive chemicals and herbivores overcoming them by detoxification or sequestration (Agrawal et al., 2009; Zangerl and Berenbaum, 2005). We also have data from parasitic mistletoes and their hosts, a case which may be more likely to generate inverse matching dynamics expected for parasites resisted by host immune recognition (van Halder et al., 2019; Yan, 1993). Among the mutualistic interactions represented in our data, we have some indirect evidence for host-mutualist matching dynamics (in *Yucca*/*Tegeticula*, Smith et al. 2009; in *Medicago*/*Ensifer*, Yoder 2016). However, some of these interactions may also be dominated by host selection against “cheating” genotypes in the associate species, which creates directional selection more akin to an arms race (Yoder 2016), for example in ant-plant protection interactions (Heil and McKey, 2003). It is likely that the broad differentiation we make between antagonistic and mutualistic dynamics does not have the explanatory power we could obtain with direct characterization of selection dynamics in individual plant-associate pairs — though of course rigorously characterizing the selection created by species interactions is a longterm, ongoing project for the entire field of coevolution studies.

Whether a host-associate interaction will create isolation-by-host for the associate species is likely determined by the genetic basis of the host and associate traits mediating the interaction, as well as the forms of coevolutionary selection acting on those traits. For the host-associate pairs in our compiled dataset, the specific genetic basis of interacting traits is even less well characterized than the forms of selection that may be operating. Strong selection acting on a trait determined by one or a few loci is expected to create patterns of differentiation at those loci that would deviate from genome-wide population structure (Hoban et al., 2016), and this should mean that a host’s genome-wide population structure will poorly predict the variation in host-mediated selection experienced by an associate. On the one hand, simple genetic bases have been identified in classic studies of plant-parasite interactions (Flor, 1956), and in plant resistance to herbivory (e.g., Johnson et al. 2018), and in pollinator-attracting floral color and scent (e.g., Gates et al. 2018; reviewed by Grotewold 2006; Sheehan et al. 2012). However, there is also evidence for quantitative and continuously varying traits mediating plants’ resistance to parasites (Pilet-Nayel et al., 2017), defenses against herbivory (Agrawal, 2007; Fox, 1981), as well as attraction and rewarding of pollinators (Caruso et al., 2010; Fournier-Level et al., 2009; Galliot et al., 2006) — and if interactions are mediated by multiple traits, host-associate compatibility and local co-adaptation may be effectively polygenic even if individual traits are Mendelian. Thus, we think it may not be unusual to find host-symbiont compatibility broadly aligned with hosts’ genome-wide population structure.

### Opportunities for future work

The results of our literature search reveal considerable opportunities for future studies of population genetic patterns across trophic levels. Although population genetic tools have been applied to the broad subject of plant-associate diversification for decades, we find just 15 papers reporting data for both a plant and at least one intimate associate species — 11 published in the last 10 years. The majority of these studies use codominant elecrophoretic markers, which have relatively poor resolution to characterize diversity and differentiation compared to modern DNA sequencing methods; and this is reflected in the wide range of precision we find when we estimate correlations between genetic, geographic, and environmental distances (Fig 2). We expect that future studies of this type will benefit greatly from the increased accessibility of genome-wide sequence data (Andrews et al., 2016; Davey et al., 2011).

Finally, 16 of 20 associate taxa in our compiled data set are insects. This pattern is not surprising in light of the historical focus on angiosperm-insect co-diversification (Ehrlich and Raven, 1964; Farrell, 1998; Farrell et al., 1992; Futuyma and Peterson, 1985; Peterson and Denno, 1998). However, many of the same reasons to expect that insect herbivores and mutualists should evolve in response to their host plants apply to symbiotic fungi (e.g., Escudero 2015), bacteria (Harrison et al., 2017), and even parasitic or commensal plants (Jerome and Ford, 2002; Schneider et al., 2016; Walters et al., 2021) — other cases in which a single associate individual lives most of its life on one host, and in which associates’ geographic dispersal is limited by hosts’ distribution. Moreover, plants are far from alone in hosting parasitic and mutualistic associates in intimate interactions. It is easy to imagine a version of the present analysis examining animal hosts and microbial symbionts, parasitic nematodes, or pathogenic viruses. One of the clearest opportunities for further contributions to this line of inquiry is the expansion of its taxonomic scope. Broadening the range of species associations represented with this kind of data will further open the possibility of comparing outcomes for interactions with varying degrees of specificity — we would expect that obligate, specialized associates will be more likely to experience ecological isolation arising from their host’s population structure, but this is not testable with the limited range of species represented in our current data set.

### Conclusions

Compiling population genetic data for plants and closely associated taxa, we find support for the widespread understanding that intimate interactions across trophic levels create diversifying natural selection. Although both antagonistic and mutualistic associates frequently evolve population structures that parallel those of their host plants, our analysis finds that antagonists are somewhat more likely than mutualists to experience ecological isolation attributable to host population structure. Future studies of plant-associate population genetics will, we hope, expand the taxonomic scope of this data, to further illuminate how plant-associate interactions have fueled the diversification of life on Earth.

## Acknowledgements

We thank Christopher I. Smith, John N. Thompson, and three anonymous reviewers for helpful comments on earlier drafts of this paper. Support was provided by start-up funds from California State University Northridge and the U.S. National Science Foundation (DEB, 2001180).

## Author contributions

All authors contributed to the literature search and data compilation; JBY conducted data analysis; all authors contributed to drafting the manuscript. All authors reviewed and approved the final text.

## Data accessibility

Supporting data and scripts have been uploaded to the Dryad repository, doi 10.5061/dryad.fxpnvx0w3.

